# Controlling and measuring dynamic odorant stimuli in the laboratory

**DOI:** 10.1101/733055

**Authors:** Srinivas Gorur-Shandilya, Carlotta Martelli, Mahmut Demir, Thierry Emonet

**Affiliations:** Interdepartmental Neuroscience Program, Yale University; Department of Molecular, Cellular, and Developmental Biology, Yale University, New Haven CT 06511; Department of Physics, Yale University, New Haven, CT 06511

**Author notes:** Volen Center for Complex Systems, Brandeis University, Waltham MA 02453. Department of Biology, University of Konstanz, Konstanz, 78457 Germany (Carlotta Martelli).

## Abstract

Animals experience complex odorant stimuli that vary widely in composition, intensity and temporal properties. However, stimuli used to study olfaction in the laboratory are much simpler. This mismatch arises from the challenges in measuring and controlling them precisely and accurately. Even simple pulses can have diverse kinetics that depend on their molecular identity. Here, we introduce a model to describe how stimulus kinetics depend on the molecular identity of the odorant and the geometry of the delivery system. We describe methods to deliver dynamic odorant stimuli of several types, including broadly distributed stimuli that reproduce some of the statistics of naturalistic plumes, in a reproducible and precise manner. Finally, we introduce a method to calibrate a Photo-Ionization Detector to any odorant it can detect, using no additional components. Our approaches are affordable and flexible and can be used to advance our understanding of how olfactory neurons encode real-world odor signals.

## INTRODUCTION

The study of sensory systems requires precise control and measurement of the stimulus applied. The ease of generation and measurement of light and sound stimuli have led to a detailed understanding of how primary visual and auditory neurons encode visual and auditory stimuli^1–4^. Olfactory stimuli are harder to generate and measure because they consist of small molecules of bewildering variety that have to be transported from the source to Olfactory Receptor Neurons (ORNs) on sensory organs.

The kinetics of odor stimuli depend on both physical and chemical parameters^5,6^. Far from surfaces, odors are transported by advection and diffusion. Because these processes depend linearly on odor concentration, the kinetics of odor transport are independent of odor concentration. This means that, if interactions with surfaces could be avoided completely, odor pulses of different concentrations could in principle be delivered with exactly the same kinetics. In practice, surfaces are unavoidable and close to them, odor molecules can diffuse through the flowing air to the surfaces where they can bind. The binding of odor molecules to surfaces is a nonlinear process because it depends on the concentration of odor in the gas-phase and that of odor molecules already bound to the surface. Thus, surface interactions break the linearity of odor transport, which renders the kinetics of odor transport *concentration-dependent* (See Methods). Natural odors can be composed of multiple odorants with different surface affinities. Surface interactions might affect some components of an odor more than others, possibly breaking the coherence of an odor signal. These interactions can significantly alter the kinetics of identically delivered stimuli, complicating the analysis of ORNs responses^6,7^. Surface-odorant interactions are significant both in laboratory conditions used for physiology and behavior^8–15^, and in real-world scenarios^16–20^.

In this paper, we aim to provide simple approaches to make the delivery of complex odorant stimuli more controllable and reproducible. We first analyze the properties of a standard odor delivery system using a simple mathematical model. We aim to provide some intuition about why the output of the delivery system may not match expectations. Previous works have used either simple odor delivery systems that can generate simple stimuli, or custom-designed instruments capable of complex stimuli but that are challenging to engineer^6,21-24^. Here, we describe methods to deliver complex and intermittent odorant stimuli using off-the-shelf components. We show how the same delivery system can be used to reproducibly generate an odorant stimulus that mimics the statistics of natural stimuli. Finally, we describe a novel method to calibrate a Photo-Ionization Detector (PID) to any odorant it can detect, allowing researchers to relate physiological and behavioral responses to absolute odor concentrations.

## MATERIALS AND METHODS

### Odor delivery system

The odor delivery system shown in Figs. 2a, 3a is built of a few off-the-shelf components that can be reconfigured based on the task. The key ingredients are as follows: a source of pressurized air to power the system and generate flows, instruments to regulate flows (Mass Flow Controllers or MFCs), valves to divert flows, tubing to connect different parts together, a glass tube to deliver the odorized airstream to the preparation, and a device to measure the output (a Photo-Ionization Detector, or PID). For a source of pressurized air, we used “dry air” from Airgas (item # AI D300) which is purified compressed atmospheric air. Air from this pressurized cylinder was delivered to a set of MFCs (Alicat Scientific MC-series) that regulated flow to downstream components. We chose these MFCs since they can be interfaced with in a number of ways, including via USB, and provide programmable parameters of the underlying control system, allowing us to tune MFCs to desired applications (for example, a MFC that is dedicated to maintaining a steady flow that does not change can be configured with control parameters that enhance stability at the cost of speed, while a MFC that is used for rapidly varying flow through a odorant vial to generate dynamical odorant signals can be optimized for speed of response). We used solenoid valves (Lee Co, LHDA 1231515H) to divert flows either to the preparation or to waste. Pure odorant was contained in generic 30mL glass scintillation vials. We drilled holes in the cap and threaded them so we could fit them with push-to-connect tube fittings (McMaster Co, 1/8: tubing, 10-32 UNF 52065K211). This allowed for a modular container to hold pure odorants that could quickly be connected to the rest of the system.

To connect different components together, we used 1/8” Teflon tubing (McMaster Carr #5239K24) since odorants stuck to this material minimally (other common tubing materials such as soft Tygon plastic bound far more to common laboratory odorants, and should be avoided). A good test of the material being used for tubing is to deliver an odorant pulse through it, disconnect the source, and deliver another pulse. If odorants bind to the material extensively, the second control pulse will not be entirely flat. To connect tubes to each other, we used push-to-connect fittings (e.g. McMaster Carr #5111K102, #5779K21, #5779K41).

No component is perfect, and it is important to understand the limitations of the parts being used to build an odor delivery system. Neglecting these shortcomings can lead to unanticipated and hard-to-diagnose failure modes in the assembled system. For example, for the Lee valves we used, their series resistance varied substantially from valve to valve, meaning that it was impossible to use them in parallel to build a symmetric switch. While using Teflon tubes can minimize contamination with odorants, odorants can still adhere to the interior of tubes. To minimize contamination, we used separate tubes and connectors for each different odorant. The interior of the solenoid valves we used are not coated with Teflon, and therefore we also used a different valve for every odorant.

### Software

Many of the results we present in the text were made possible by converting the problem of configuring and tuning hardware elements into a software optimization problem. This enabled parameter searches that would be otherwise prohibitively tedious. We wrote controller (available at https://github.com/sg-s/kontroller), a MATLAB-based data acquisition and control system that we wrote and could automate many tasks. A small MATLAB toolbox that tunes the control parameters of MFCs is available at https://github.com/sg-s/alicat-MFC. Finally, software to interactively optimize control signals to deliver naturalistic odorant signals (like in Fig. 2) is available at https://github.com/sg-s/methods-paper/, which can be used as a template to build other optimization routines.

### Mathematical model of a simple delivery system

In this section, we consider a delivery system schematized in Fig. 1a. Clean air is injected at the constant flow rate Q inside a cylindrical odor delivery tube of inner radius *R*. From a lateral hole a secondary air stream carrying odor at gas phase concentration c1 is injected in the same tube with flow rate *Q*_1_.

**Figure 1.**
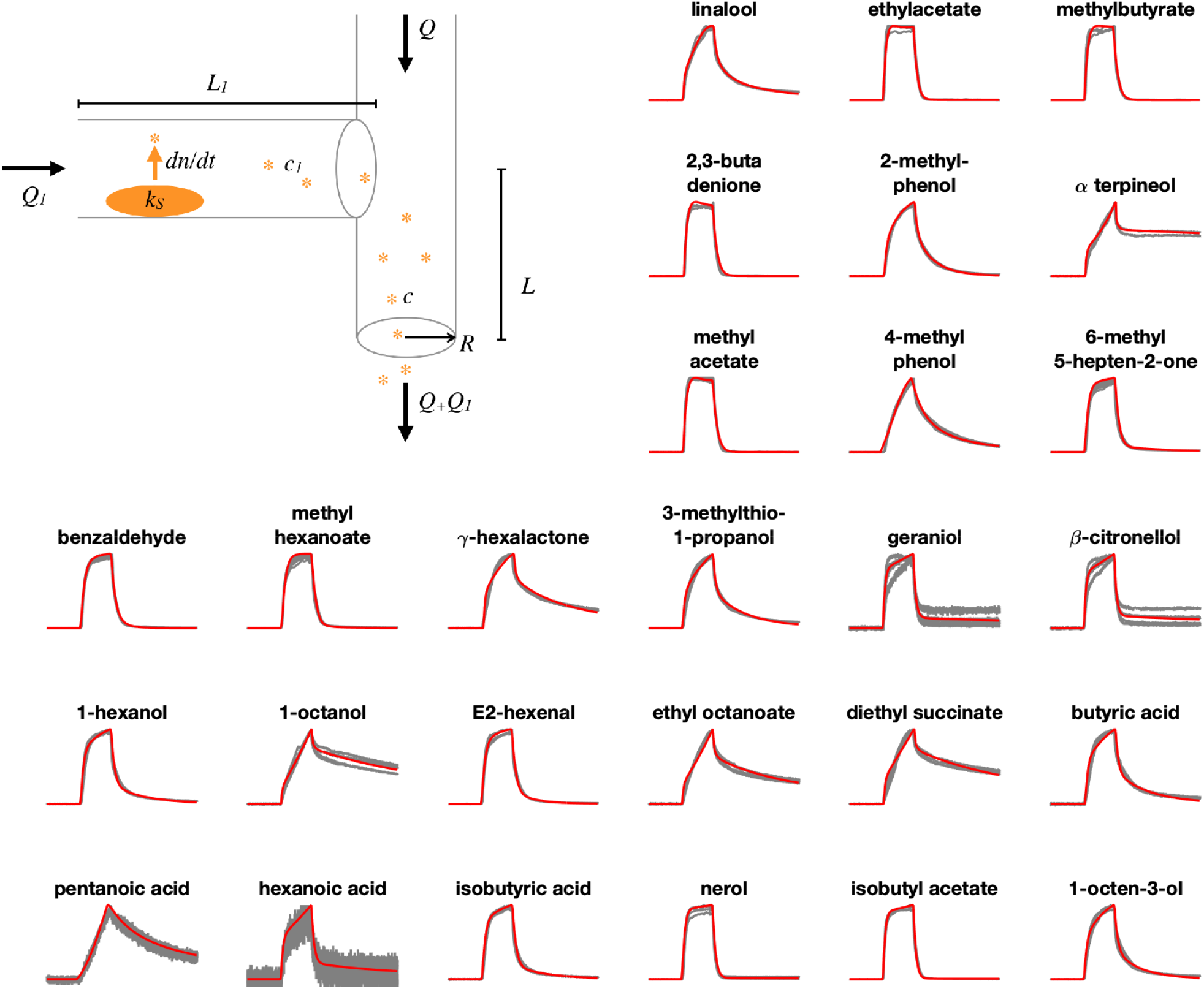
A simple model reproduces the diversity of odorant pulse kinetics. A simple method of delivering odorants is to blow air for a short duration through a chamber contained odorized air. One way to create an odorized chamber is to insert a small piece of paper soaked in liquid odorant. Odorants delivered this way exhibit a broad range of stimulus kinetics, that depend on the chemical identity of the odorant. Each panel shows five measurements of the kinetics of stimulus using a PID, normalized by the peak (black traces). Our model reproduces the observed diversity in odorant pulse kinetics: in each panel, red curves are the model fits (Eq. [5-6]). The pulse duration is 500 ms. Data here is from Fig. 1b in ^6^.

**Figure 2.**
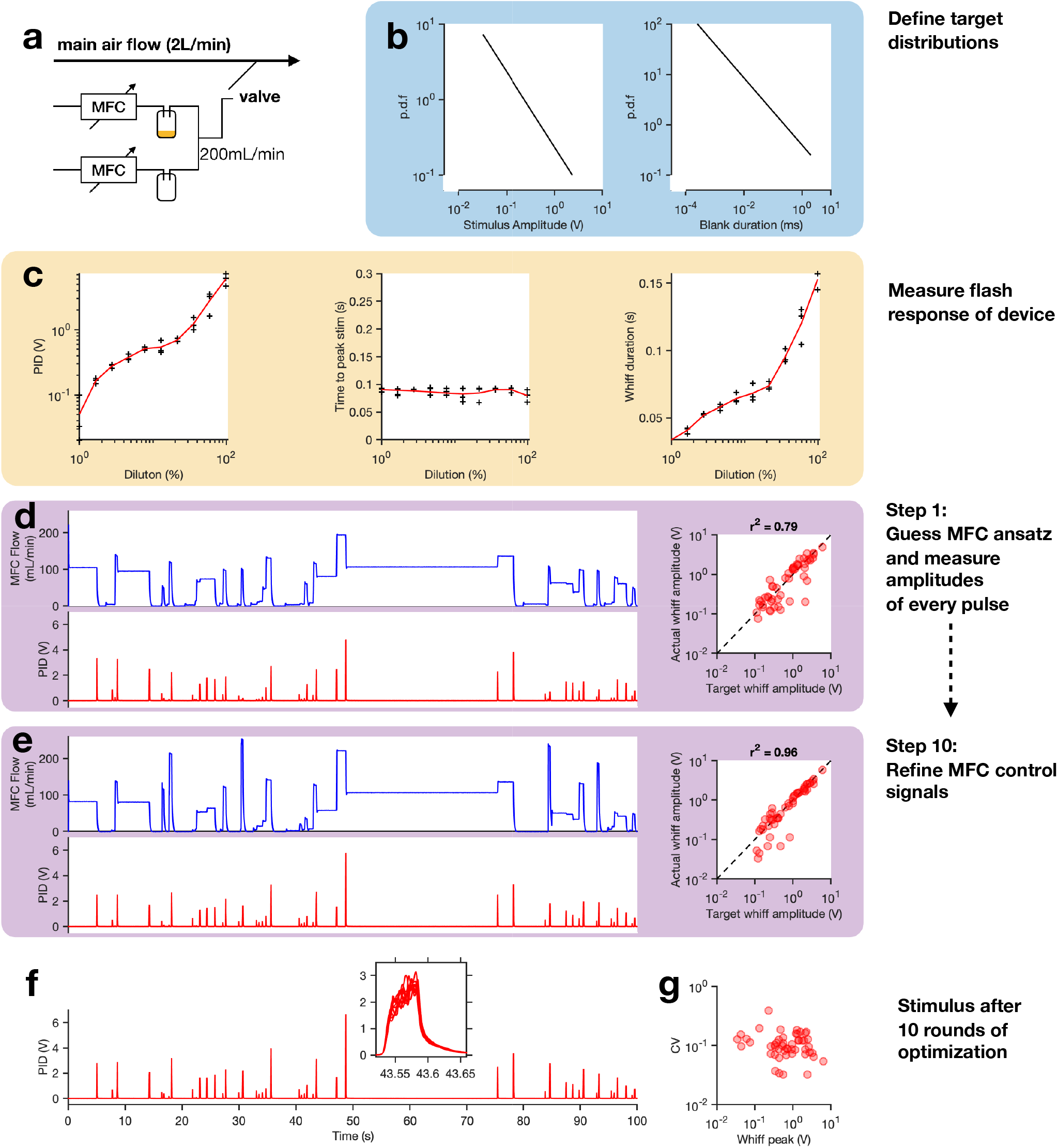
How to deliver odorant stimuli with naturalistic statistics. (a) A delivery system consisting of two MFCs is used to vary the dilution of an odorized airstream in a main airstream. (b) Target distributions of whiff amplitude and blank duration. (c) Measuring the flash response of the device. Using the flash response data, an ansatz MFC signal and sequence of valve opening can be constructed. (d) Measurement of odorant concentration in response to this MFC ansatz. Correlation between measured whiff amplitudes and desired whiff amplitudes. Based on measured mismatches between the amplitude and duration of every whiff, the control signal to the MFC can be iteratively optimized. (e) Odorant signal after ten rounds of optimization. Whiff amplitudes are now close to the target whiff amplitudes, for every whiff. (f) The final stimulus matches the target statistics, and is highly reproducible (inset). (g) Coefficient of variation of whiff amplitude vs. whiff peak, for all whiffs, showing no significant trend.

In the absence of any interactions between odorant molecules and surfaces, the gas phase concentration of odor inside the delivery tube, *c*, obeys the equation

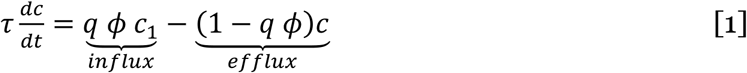

The first and second terms on the right represent the odor that comes in and out of the delivery tube, respectively. τ = *V*/*Q* is the time it takes to replace all the air in the delivery tube (*V* is the volume of the tube downstream from the lateral inlet) and ϕ = *Q*_1_/*Q* is the ratio of the flow rates injected in the two tubes, which in experiments is typically less than 1. We assume that the odorized air injected through the lateral hole at concentration *c*_1_ mixes rapidly with the clean air stream. *q* is the state of the valve that controls the odorized air stream: before the pulse *q* = 0 and during the pulse *q* = 1. Equation 1 is linear in the concentration *c*. Thus, if we double the concentration at the input, *c*_1_, the concentration at the output *c* will also double. Nothing in this equation depends on the identity of the odor, indicating that in the absence of interactions with surfaces, the time-dependent shape of the odor pulse delivered is independent of the identity of the odorant.

In general odorant molecules always interact with the surface of the delivery tube. To account for the loss of odorant molecules from the gas phase to the inner surface of the tube we add an extra term to the equation:

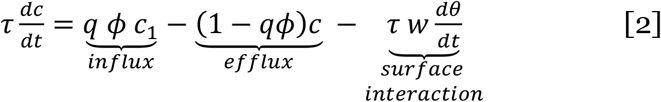

here 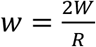, where *W* represents the surface density of binding sites for odorant molecules. This equation must be supplemented with an additional equation that describes how the fraction, θ, of the tube surface that is covered by odorant molecules, changes as a function of time:

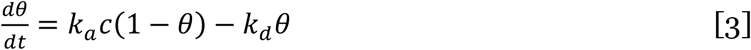

Here ka and kd are the binding and unbinding rates. To gain intuition about the effect of surface interactions on the stimulus dynamics it is instructive to make the simplifying assumption that the odorant-surface interactions are much faster than the transport of odorant by the flow. Solving Equation 3 at quasi-steady state and inserting in Equation 2 yields

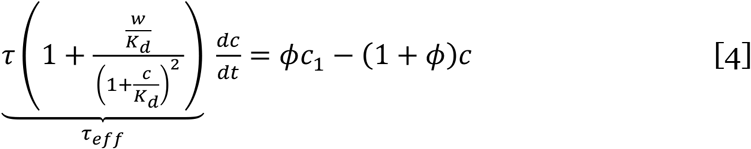

where *K*_*d*_ = *k*_*d*_/*k*_*a*_. Comparing this with Equation 1, we see that the effective timescale τ_*eff*_ of the stimulus kinetics now depends on several parameters of the odor-delivery system and on the odorant concentration. Equation 4 indicates that the effect of surface interactions becomes significant when the effective concentration of surface binding sites per volume w=2*W*/*R* becomes much larger than the dissociation constant *K*_*d*_. Thus, one way to mitigate such effects is to use a tube material that is more inert therefore reducing *W* and increasing *K*_*d*_. One can also reduce *w* by increasing the inner radius *R* of the delivery tube. However, this is costly and often not practical because an increase in the radii leads to a quadratic increase in the volume of air and odor that must be flown through the delivery system. For odorants that do not interact extensively with surface, i.e. when *c* ≪ *K*_*d*_, the denominator on the left-hand side of the equation becomes 1 and Equation 4 becomes linear in *c*. In this case, the kinetics of odor pulse are independent of the odor concentration. Odorant that satisfy these conditions (*w* ≪ *K*_*d*_ or *c* ≪ *K*_*d*_), such as ethyl acetate, were called “fast” odorants in ^6^. In contrast, for odorants that interact significantly with surfaces, i.e. when *c* is on the same order as *K*_*d*_, Equation 4 predicts that the pulse kinetics depend on odor concentration. As the odor concentration becomes smaller, the time scale τ_*eff*_ becomes larger. These odorants were called “slow” in ^6^.

To complete our simple model of the delivery system in Fig. 1 we must also model the odor concentration in the gas phase, *c*_1_, and on the surface, θ_1_, of the tube used to inject the odor into the delivery tube. Similar to Equations 2 and 3 we have:

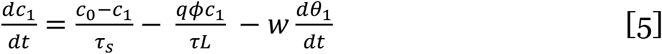

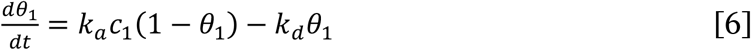

where *L* = *V*_1_/*V* is a geometric factor that depends on the relative length of the two tubes if their radius is the same. The first term on the right-hand side of Equation 5 represents the dynamics of equilibration between the odor concentration right above the liquid source, *c*_0_, and the gas phase concentration in the tube, *c*_1_. The time scale, τ_*s*_, depends on the diffusion coefficient of odorant molecules in the gas phase and the air dynamics in the head space. When using pure odorants in liquid phase and when the system allowed to reach equilibrium, *c*_0_ is simply the saturated vapor pressure. When odors are diluted in paraffin oil, or when multiple odorants are mixed in the liquid phase, *c*_0_ becomes a complex function of the concentrations of odorants in the liquid phase because the presence of one compound can affect the evaporation rate of another one. These complex effects are manipulated by the perfume industry to control the timing of release of various “notes” of a perfume^25^.

After normalizing *c* and *c*_1_with the concentration *c*_0_ Equations 2, 3, 5 and 6 provide a set of 4 ordinary differential equations that we can use to fit pulse kinetics of different monomolecular odorants (see **Results and Discussion**). Except for the 4 parameters τ_*s*_, *k*_*a*_*c*_0_, *k*_*d*_, *w*/*c*_0_, that depend on odorant identity, all other parameters are known.

### How to estimating the dose-response curve of an olfactory neuron using odorant pulses

Short pulses of odorants are usually used to estimate the dose-response curve of single neurons^6,24,26^. A simple way to generate odorant pulses of varying concentrations is to force air through cartridges containing odorants diluted to varying degrees in a solvent, like paraffin oil. Supplementary Fig. 1a shows a typical dose-response measured by delivering odorants using this method. The peak firing rate of the ORN being measured from, ab3A, varies non-linearly with the liquid phase concentration of the odorant in the cartridge and can be fit by a Hill function. PID measurements of the stimulus intensity for every trial allow us to compare measured stimulus intensity to the nominal liquid phase concentration (Supplementary Fig. 1b). Significant deviations from linearity and considerable trial-to-trial variability are evident (Supplementary Fig. 1b). Moreover, plotting the response of ab3A against the measured gas-phase concentration (Supplementary Fig. 1c) yields different parameters from a fit with a Hill function. This suggest that the liquid-phase concentration is not always an accurate proxy for the gas-phase concentration and can thus bias estimations of response properties of neurons.

Most trial-to-trial variability in the stimulus is due to a slow decay in the odor concentration, as shown for four odorants at different liquid phase concentrations (Supplementary Fig. 1d-h). This problem is mostly due to the depletion of the liquid dilution which depends on the volatility of the compound used (cf. ethyl acetate vs. diethyl succinate)^5^.

One way to ameliorate this problem is to use a Mass Flow Controller (MFC) to vary the airflow through a scintillation vial containing pure odorants (Supplementary Fig. 1i-m)^8,23,27,28^. The concentration of odorant in the odorized airstream depends only on the flow rate of the MFC and it is stable over time as far as there is some odorant left in the vial and the air is blown continuously through it (and sent to exhaust). In this configuration, the solenoid valve is then placed downstream of the odor source. Pulses delivered using this method are more “square” than the previous method (Supplementary Fig. 1d, i), and do not show any significant decay in amplitude over 10 trials (Supplementary Fig. 1j-m), even for volatile odorants like ethyl acetate. Using this method allows the experimenter to easily switch between different stimulus amplitudes by changing the flow rate, allowing automation of data collection.

### How to deliver intermittent odorant signals

Odor cues seldom occur as isolated pulses in natural settings. Instead, they arrive in intermittent series of pulses whose statistics carry information about the odor source location^18^. Thus, various types of fluctuating odorant stimuli have been used to characterize the dynamic response properties of ORN^6,22,23,29,30^. As in other sensory modalities^31,32^, paired recordings of a fluctuating sensory signal and the corresponding neural response have be used to estimate the linear kernel that best describes the transformation from the stimulus to the response^6,22,23,33,34^.

The linear kernel can then be used to predict the time series of the response of that neuron to a novel stimulus that the neuron has not previously been exposed to. For example, this approach revealed that for many odor-receptor combinations in the *Drosophila melanogaster* antenna, the linear response kernels of ORNs were more invariant than previously anticipated^6,35^. A favorable stimulus to use for this purpose is Gaussian white noise. Its tight autocorrelation structure allows sampling as many frequencies of the neuron response as possible. Using such a stimulus, linear filters can be estimated in an unbiased fashion even in the presence of an output nonlinearity^31^.

However, Gaussian white noise odorant signals are hard to realize. Interaction of the odorant with the walls of the delivery system^6^, limits on the airspeed used in the delivery system, and non-infinitesimal timescales of the components of the delivery system introduce temporal correlations into every odorant stimulus. Hence in practice the stimulus is never “white”. Furthermore, since the easiest way to control an odorant signal is to use a valve to divert an odorized airstream towards or away from the preparation (animal subject), early work using fluctuating odorant signals to identify linear kernels from ORN responses used binary odorant stimuli where the stimulus was either on or off^6,22,36-39^ (Supplementary Fig. 2a-b). This tends to generate a bimodal stimulus distribution (Supplementary Fig. 2e-h), with the lower peak close to zero, and the larger peak at some value that can be controlled by the concentration of the odorized airstream (Supplementary Fig. 2a-b). The correlation time of the signal can be controlled by varying the switching time of the valve, and for volatile odorants that do not interact strongly with the surfaces of the delivery system, correlation times as fast as 30ms can be realized^6^ (Supplementary Fig. 2i-l). In these binary stimuli, the mean stimulus is typically rarely realized, and the responses of the neuron are dominated by responses to the large, rapid increases of odorant on valve opening^6,22^.

An alternative approach can deliver intermittent odorant signals that have a unimodal, more Gaussian distribution. Using a MFC to vary the flow rate of an airstream through a vial of pure odorant generates an odorized airstream with a gas phase concentration that depends on the flow rate through the odorant vial (Supplementary Fig. 2c-d)^23,29^. Odorant stimuli delivered this way typically don’t need to “bottom out” at zero-stimulus (Supplementary Fig. 2e-h), and have mean values close to their mode. The autocorrelation time of a stimulus delivered this way is limited by the update rate of the MFC (Supplementary Fig. 2i-l). If odorant stimuli with tighter autocorrelation functions are desired, multiple MFCs can be chained in parallel, driven by uncorrelated control signals (not shown).

## RESULTS AND DISCUSSION

### Basic parameters that affect the kinetics of an odor pulse

A common method of delivering odorants is to create a chamber of odorized air by evaporation of liquid odorant (Fig. 1) and push that odorized air into a main delivery tube where a constant stream of clean air is flowing. Signal intensity can be controlled by diluting the odor in liquid phase (e.g. mixing pure odor with paraffin oil) ^21,40,41^. A solenoid valve upstream of the odorant controls odor delivery timing while minimizing contaminations. This system has been used to screen through large numbers of odorants^10,42^ and to characterize the sensitivity of Drosophila ORNs to ecologically relevant stimuli^10,21^. Using this approach to measure the response kinetics of ORNs, in particular the linear response function of neurons, has been more challenging. PID measurements have revealed that stimuli delivered in this way typically exhibit intrinsic odorant-dependent dynamics before any interaction with the animal takes place^6^ (Fig. 1) which therefore can obscure the neural response function of ORNs.

To gain intuition about the parameters that affect odor stimulus dynamics we created a simple mathematical model of the delivery system schematized in Fig. 1 (see **Materials and Methods**). We fitted this model to pulse kinetics of 27 different monomolecular odorants commonly used in the laboratory (Fig. 1, red lines). We find that the dynamics of odor pulse vary according to odor identity due to differences in the following four parameters: the characteristic time scale of liquid to gas transition at the odor source, the rate of binding and unbinding of the odor to the surfaces of the delivery tube, and the amount of odorant molecules that can bind the surface before it becomes saturated. The model successfully reproduces the diversity of observed kinetics of odorant pulses, including relatively sharp pulses (e.g. Fig. 1, ethyl acetate), slow kinetics (e.g., Fig. 1, 2-methyl phenol) and super-slow decays following pulse offset (e.g. Fig. 1, α-terpineol). This model can be easily fit to experimental data from other delivery systems, and can guide intuition on the feasibility of using certain odorants with specific dynamic stimulus patterns. For a full description of how different parameters influence pulse kinetics see **Materials and Methods**.

### Delivering odor stimuli with naturalistic and defined statistics

Natural odor signals are comprised of whiffs and “blanks” whose intensities and durations can be distributed over many orders of magnitude^18,20,43^. A fan can be used to drive turbulent air over an odor source, mimicking natural plumes, but the resulting stimulus pattern is not reproducible^6,12,22,44^.

Our approach is to deliver an odorant signal whose whiff and blank statistics are drawn from a known distribution. We exploit the fact that some naturalistic signals can be approximately segmented into short pulses. Therefore, a target signal can be approximated by adjusting the amplitude, duration and intermittency of a set of single whiffs. The experimental apparatus we use is shown in Fig. 2a. A pair of MFCs regulate airflow through an odor vial, controlling odor concentration. A valve enables pulse timing control. The whole system can be modelled as a mapping from MFC and valve control signals to the measured whiff statistics. Thus, control signals can be iteratively tuned until the measured signal is sufficiently close to the target signal.

First, we define target distributions (power laws) of whiff intensity and the durations between whiffs (Fig. 2b). We first characterize the flash response of the odor delivery system, which determines the control signals to the MFC and valve needed to generate a pulse of a desired amplitude. Setting MFC flow rates to some fixed value, we briefly activate the valve, and measure the resultant PID waveform. By repeating this procedure over multiple MFC flow rates, we can fit functions that approximately describe how the pulse statistics depend on the dilution (Fig. 2c). We then generate a target stimulus time-series by repeatedly sampling from the distributions in Fig. 2b. We first sample a whiff intensity, then draw another sample from the blank duration distribution, and so on, till a skeleton of the odor stimulus is created. Since we sample from the target distribution in Fig. 2b, we are guaranteed that these statistics match the target distributions.

Next, we generate an ansatz control signal for the MFC and for activation of the valves, by inverting the functions fit in Fig. 2c. The MFC ansatz is shown in Fig. 2d (blue trace). Measuring the stimulus using the ansatz control sequence reveals a sequence of whiffs with varying amplitudes, but their amplitudes are not perfectly correlated (*r*^2^ = 0.79) with desired whiff amplitudes (Fig. 2e). These deviations stem from the fact that mapping in Fig. 2c assumed isolated flashes, which is not true in this stimulus. We therefore built a simple optimizer that varies the set point of the MFCs before each valve opening till the correlation between measured whiff intensities and desired whiff intensities is within acceptable limits. Ten rounds of this optimization increased the correlation sufficiently (*r*^2^ = 0.96) (Fig. 2e).

Once a desired accuracy has been reached, the stimulus can be replayed without further fine-tuning. Fig. 2f shows ten repetitions of the same stimulus, showing high reproducibility despite the nearly 1000-fold variation in signal intensity across whiffs. The coefficient of variability (CV) for each whiff is well-controlled, and does not depend systematically on whiff intensity (Fig. 2g). Similar strategies can be used to generate odorant signals with defined means and variances (Supplementary Fig. 3).

### Calibrating PIDs

The PID has become a common tool to measure odorant stimuli, since they are fast and easy to use. PIDs continuously sample air and return a voltage value proportional to the number of ionizable molecules in the air sample, but do not report an absolute value of the gas phase concentration of odorants. Studies that focused on characterizing temporal features of the stimulus do not necessarily require a calibration of the PID ^6,22,45^. However, a calibration is necessary when one wants to compare the sensitivity of a receptor to different odorants^5^, or to compare stimuli, physiological and behavioral responses across laboratories.

The PID can be calibrated by coupling the PID with another device that can measure absolute concentrations, like GCMS devices or flame ionization detectors^5,46^. However, this shifts the problem of calibration to another system, that in turn needs to be calibrated. Another approach is to assume the odorant headspace in the olfactometer is saturated, and use Raoult’s Law and Henry’s Law to estimate the gas phase concentration of the odorant, and then extrapolate to fast changes reported by the PID. Such an approach has been used in ^47,48^ or in conjunction with a tracer gas of known concentration^23,29,36,37^. It is not known how valid the assumption of saturated headspace is, and using another tracer gases once again moves the problem of calibration onto another system.

Here, we propose a simple method to calibrate the PID to any odorant that it can detect. This method does not require additional equipment or gases of known concentration, nor does it make assumptions about the saturation of headspace in the odor container. It works by depleting a known volume of pure odorant, and integrating the total PID signal over the course of depletion. Since the number of molecules in the pure odorant sample can be estimated precisely using micropipettes and known molecular weight and density, the PID signal per unit time can be converted into the number of molecules leaving the delivery tube per unit time.

To calibrate our PID to ethyl acetate, we placed 100 µL of pure odorant in a 30mL scintillation vial downstream of a MFC, as shown in Fig. 3a. We forced air through the scintillation vial till all odorant was depleted and repeated this for a few fixed flow rates. The volume of air and time required to evaporate all odorant decreased with the flow rate (Fig. 3b-c). PID signal amplitudes increased with flow rate. If the PID captured all the odorant molecules, and ionized all of them, the integrated PID signal would be a constant, corresponding to the total number of molecules of odorant in the vial. However, we observed that the total integrated PID signal decreased with the flow rate (Fig. 3d), an effect we attributed to flow-rate dependent partial capture of odorant molecules. To compensate for this, we fit an interpolant to this data (Fig. 3d, red line), and used this to correct for variations in the total signal. Integrating the corrected PID curves yielded curves of cumulative odorant vs. time that reached approximately the same height, corresponding to the calculated number of moles of odorant (dashed line, Fig. 3e).

**Figure 3.**
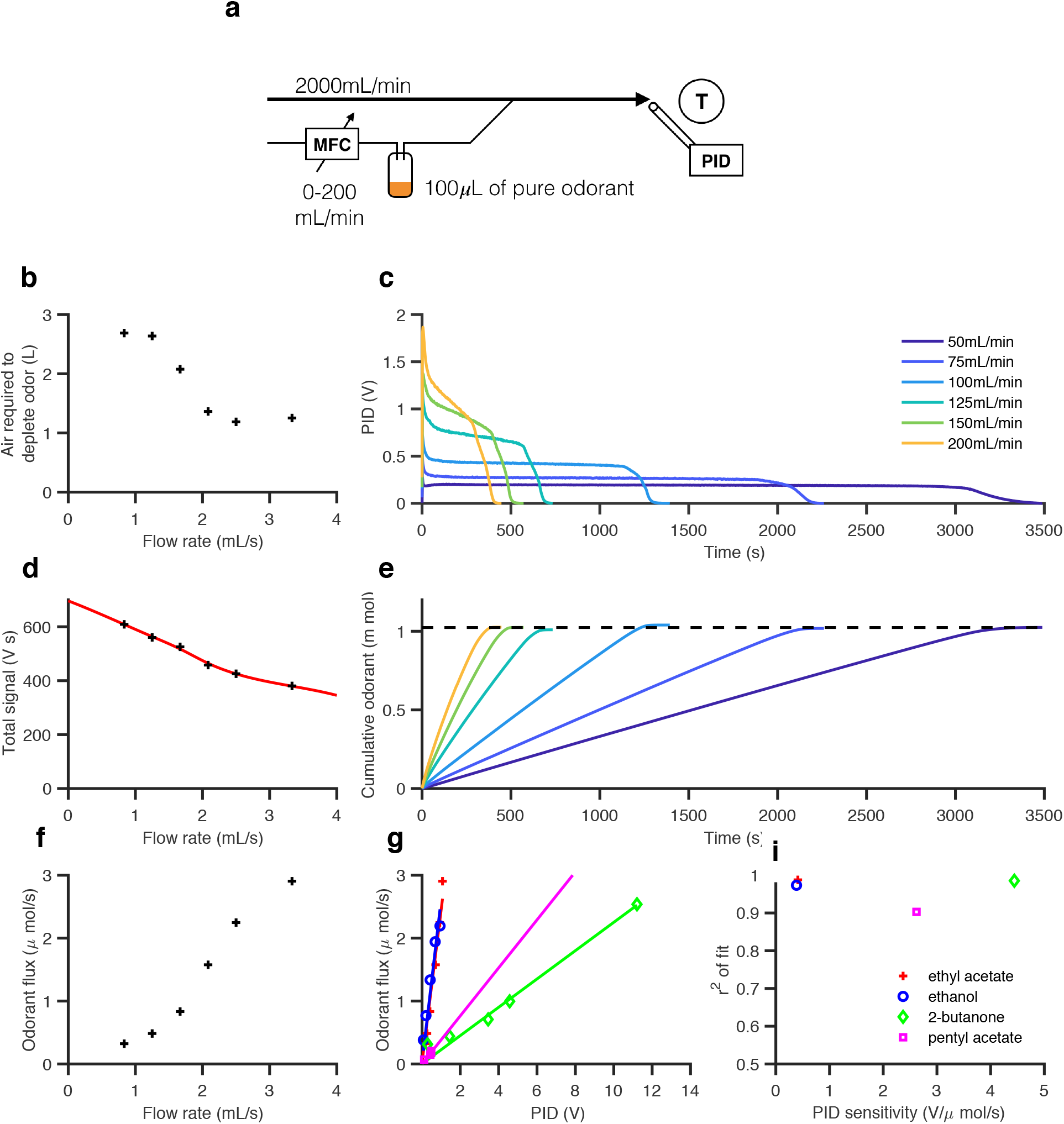
How to calibrate a PID to any odorant it can detect. (a) Schematic for PID calibration. 100 µL of a pure odorant (ethyl acetate) is placed in a scintillation vial, and air is blown over it at a fixed flow rate. A PID is placed the outlet tube close to the target (T). (b) Air required to completely deplete the odorant vs. flow rate. (c) PID traces from start to complete depletion of odorant for various flow rates. (d) Integrated PID signal as a function of flow rate. The red line is a spline interpolant. (e) Cumulative odorant vs. time for various flow rates. The dashed line indicates the total amount of odorant placed in the scintillation vial. (f) Mean odorant flux as a function of flow rate (g) Odorant flux as a function of measured PID value, for ethyl acetate (red crosses), ethanol (blue circles), 2-butanone (green diamonds) and pentyl acetate (red squares). (i) r2 of fit vs. PID sensitivity for these odorants.

We then computed the odorant flux as a function of the flow rate, by estimating the slope of the cumulative odorant curves (Fig. 3f). Finally, we can combine these measurements to plot odorant flux vs. the PID signal, to generate a function that maps PID values onto odorant flux. We repeated this calibration process for three other odorants (ethanol, 2-butanone & pentyl acetate), and find that all curves are approximately linear (Fig. 3g, i).

### Hardware abstraction and automation can speed up instrument configuration

Setting up instruments and configuring them correctly is time consuming. MFCs and odor delivery systems have many degrees of freedom and many control parameters. Finding the “right” parameter set can be laborious and challenging, since changing one parameter can have unintended effects. Using a simple configuration involving one or two MFCs and a few valves, we were able to design systems that could generate a wide variety of odor signals. To do this, we first built a programmable data acquisition and control system that could control MFCs, valves, and all other components of the delivery system, and also capture data from the PID (and electrophysiology system). This meant that the problem of finding the best flow rates, valve opening times (as in Fig. 2), or other combination of parameters to generate a desired stimulus could be translated into a software problem. This let us write simple optimizers that could iteratively change control signals to MFCs and valves, since all parameters of the hardware were manipulatable via software.

### Two odor environments

Fluctuations in an odor signal are caused by moving air and therefore can carry information about the location and distance of an odor source^18,49^. How animals extract and exploit this information remains unclear. Away from surfaces, odor transport is linear, suggesting that air flow can maintain correlations between odor components, and therefore the identity of the odor. Here we illustrated (Fig. 1) how surface interactions can introduce delays in stimulus dynamics that are odorant-specific. Thus, in the presence of surfaces individual monomolecular components of an odor might experience different delays, affecting odor identity recognition^50,51^. Effectively, there are two different odor environments: close to surfaces where odors might become decomposed, and away from them where odor coherence is maintained. To what extent animals are aware of this difference and whether they consider surface proximity when interpreting odor signals remains an open question.

---

**Supplementary Figure 1:**
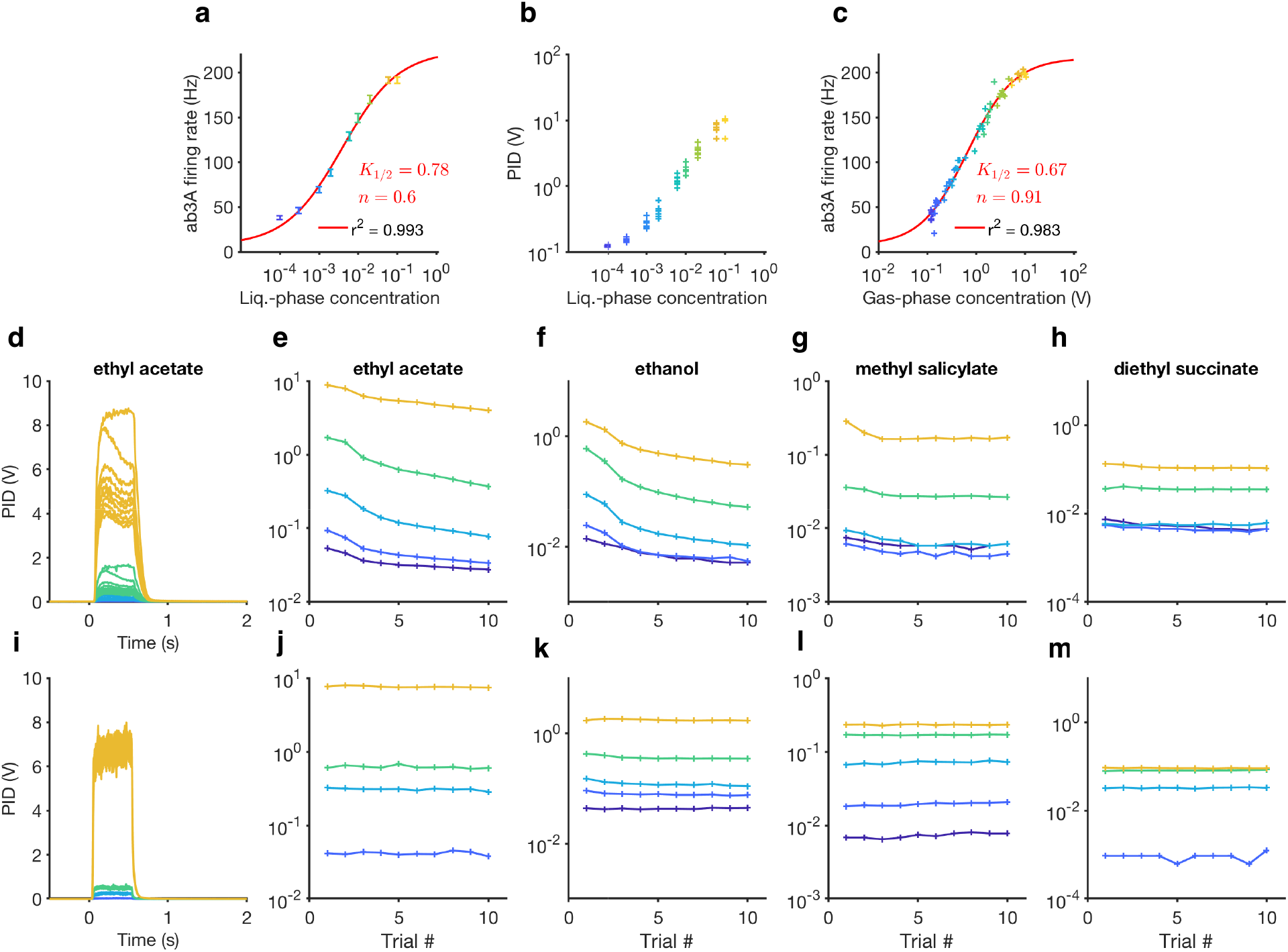
Gas-phase dilution enabled steady delivery of short odorant pulses. **(a)** Peak firing rate of ab3A ORNs as a function of liquid-phase dilution of ethyl acetate odorant in paraffin oil in “cartridges”. The red line is a best-fit Hill function. **(b)** Measured pulse amplitude *vs.* liquid phase concentration shows trial-to-trial variability and nonlinearity of measured gas phase concentration. **(c)** Peak firing rate of ab3A ORNs replotted vs. measured gas-phase concentration for every pulse. The red line is a best-fit Hill function. Note that the best-fit parameters are different in **(a)** and **(c)**. **(d-h)** 500 ms pulses of odorant delivered using liquid phase dilution. **(d)** Measured stimulus *vs*. time for different values of liquid-phase dilution of ethyl acetate. **(e-h)** Maximum pulse amplitude *vs.* trial for four different odorants. **(i-m)** 500 ms pulses of the same odorants delivered using gas-phase dilution.

**Supplementary Figure 2.**
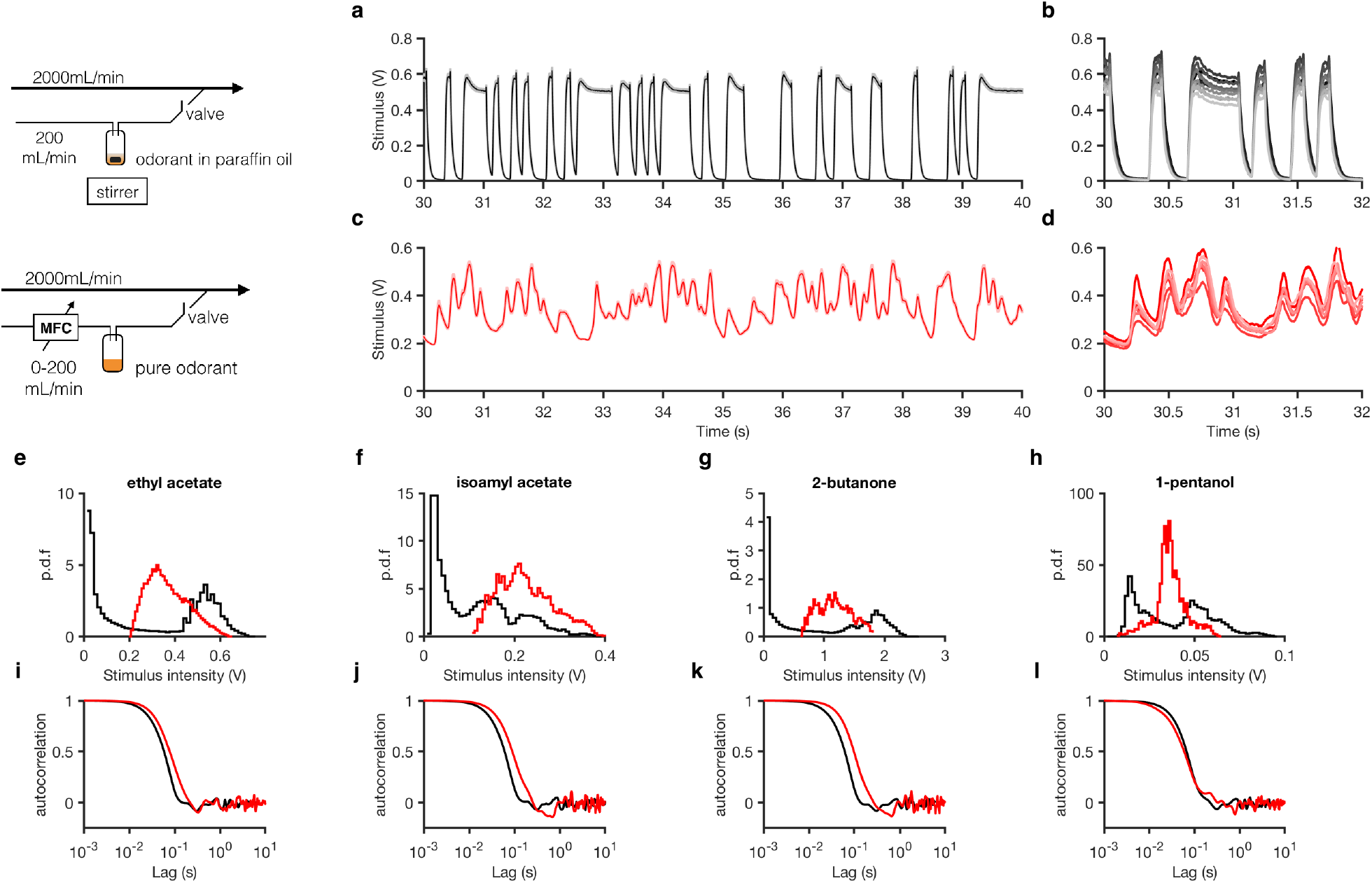
How to deliver intermittent odorant signals. **(a-b)** Using a valve to generate binary random stimuli using ethyl acetate odorant. When the valve is turned on, the odor signal tends to its maximum value, and when the valve is off, it tends to 0. **(a)** Mean ethyl acetate concentration (shading is standard error of mean). **(b)** Individual traces. **(c-d)** Using a MFC to deliver ethyl acetate with fluctuations around a desired mean. Here, the MFC is used to modulate airflow through the odorant vial, leading to fluctuations in the odor concentration. The valve is used only to shut down the airstream at the end of the odor stimulus. **(c)** Mean ethyl acetate concentration (shading is standard error of mean). **(d)** Individual traces. **(e-l)** Distributions and autocorrelation functions of stimulus time series for various odorants. Black traces are delivered using the apparatus in (a-b) and red traces are delivered using the apparatus in (c-d).

**Supplementary Figure 3:**
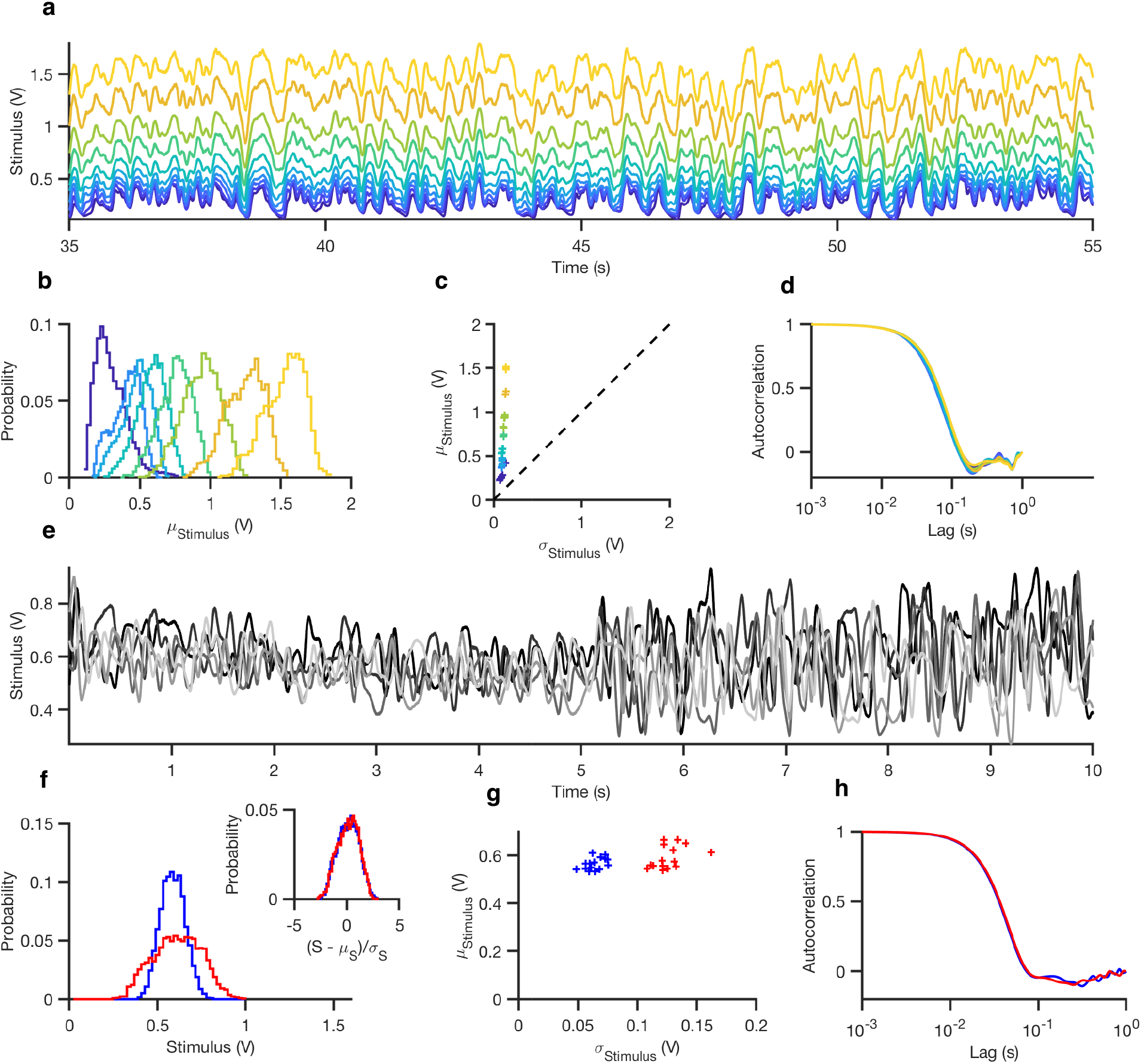
How to deliver approximately *Gaussian odorant signals with controlled means and variances*. **(a-d)** Approximately Gaussian ethyl acetate stimulus with varying mean. **(a)** Time series of stimuli with varying mean. **(b)** Probability distributions. **(c)** Mean stimulus vs. standard deviation of stimulus for each trial. **(d)** Autocorrelation functions of the stimulus for various mean values. **(e-h)** Gaussian ethyl acetate stimulus with two different variances. **(e)** Time series of the stimulus. Several trials are shown superimposed. **(f)** Distributions of the stimulus during high variance epochs (5-10 s, red) and low variance epochs (0-5 s, blue). (**f**, inset) Distributions normalized by their standard deviation, showing that they are rescaled versions of each other. **(g)** Mean stimulus vs. standard deviation of stimulus for each trial. **(h)** Autocorrelation functions of the stimulus during the high and low variance epochs.

